# Facilitating the Supplementary Motor Area Activity Reduces Variability of Ball Arrival Position in Accurate Ball Throwing Performance

**DOI:** 10.64898/2026.01.04.697346

**Authors:** Taishi Okegawa, Ayane Kusafuka, Daiki Yamasaki, Naotsugu Kaneko, Kimitaka Nakazawa

## Abstract

Maintaining high accuracy during rapid, dynamic movements is a significant challenge for the central nervous system. The supplementary motor area (SMA) is a key cortical region for orchestrating motor commands and regulating movement variability, yet its causal role in maintaining precision during high-speed tasks remains to be fully elucidated. We investigated the impact of modulating SMA excitability on throwing performance in fourteen healthy adults with no competitive throwing experience. Participants performed maximal and submaximal (50% effort) throwing tasks before and after receiving intermittent (facilitatory) or continuous (inhibitory) theta burst stimulation (TBS) over the SMA. Outcome measures included ball speed, variability of pitch location—quantified as variable error (95% confidence ellipse area) and absolute error (Euclidean distance from the ellipse center to the target)—and introspective ratings of performance via visual analog scales (VAS).

Results showed that intermittent TBS (iTBS) significantly reduced the variable error of pitch location during maximal-effort throwing—where neural noise is theoretically elevated—without compromising ball speed. In contrast, no significant changes in performance were observed following continuous TBS or during the submaximal throwing task. Notably, objective precision gains under iTBS were dissociated from subjective ratings of accuracy, which increased globally over time independent of stimulation type. These findings demonstrate that the SMA plays a critical causal role in stabilizing motor output under high motor drive. This study suggests that SMA-mediated stabilization operates as a subconscious process, offering new insights for optimizing complex motor skills.

**Highlights:**

- iTBS over the SMA significantly reduces throwing variable error in novices.
- Improved precision occurs without compromising maximal ball throwing speed.
- The SMA causally stabilizes motor output during high-speed movements.
- Objective precision gains are dissociated from subjective performance ratings.
- SMA-mediated stabilization operates as a subconscious motor process.

## Introduction

Throwing an object rapidly and accurately is a fundamental human motor skill, widely utilized in modern sports such as baseball, cricket, and handball (Roach et al., 2013). This ability emerges in early childhood even without specific training and gradually improves with ontogenetic development (Saito et al., 2025; Goodway et al., 2019). From a motor control perspective, this dynamic movement serves as an excellent model for investigating how the central nervous system (CNS) manages the trade-off between speed and accuracy. Generally, movements requiring both high speed and accuracy are subject to a speed-accuracy trade-off, making it difficult to simultaneously achieve high speed and accuracy (Fitts, 1954). A proposed physiological principle underlying this phenomenon is that the noise component of neural signals transmitting motor commands increases in proportion to the magnitude of the signal intensity (i.e., signal-dependent noise) (Faisal et al., 2008; Schmidt et al., 1979). Within this framework, skilled individuals are able to partially overcome this constraint, achieving both high speed and accuracy, whereas novices often struggle to maintain accuracy under high-speed during throwing tasks (Van Den Tillaar & Ettema, 2006). If signal-dependent noise impairs the ability to regulate variability during high-speed movements, then effective motor performance must rely on neural mechanisms that monitor and regulate the strength of motor commands to stabilize motor output.

In this context, the supplementary motor area (SMA) emerges as a key cortical structure implicated in balancing motor drive and movement variability. Anatomically located upstream of the primary motor cortex (M1), the SMA is functionally situated to orchestrate descending motor commands (Picard & Strick, 2001). Crucially, achieving high ball speed necessitates a strong neural drive, which inevitably amplifies signal-dependent noise (Gandevia, 1987). Previous studies have implicated the SMA in the generation of the “sense of effort” via corollary discharge signals, suggesting it directly monitors the intensity of this motor output (Okegawa et al., 2025; Zénon et al., 2015). Furthermore, the causal involvement of the SMA in regulating motor variability and coordination has been substantiated by non-invasive brain stimulation studies. Specifically, the disruption of SMA activity via inhibitory repetitive transcranial magnetic stimulation (rTMS) has been shown to significantly increase the variability of motor timing, underscoring the region’s essential role in maintaining temporal consistency (Jacobs et al., 2009). Conversely, increasing the SMA excitability via anodal high-definition transcranial direct current stimulation has been reported to improve the efficiency of anticipatory postural adjustments, indicating that enhanced SMA activity facilitates the precise coordination of whole-body movements (Hasui et al., 2022). Taken together, these findings suggest that the SMA functions to stabilize motor output and constrain endpoint variability. However, it remains unknown whether the SMA plays a causal role in counteracting signal-dependent noise under conditions of high motor drive, such as during maximal-effort dynamic movements.

To elucidate the causal contribution of the SMA to throwing performance, non-invasive brain stimulation offers a robust experimental approach. Specifically, theta burst stimulation (TBS), a patterned rTMS, allows for the induction of long-lasting changes in cortical excitability (Cárdenas-Morales et al., 2010; Iezzi et al., 2011). Depending on the stimulation pattern, TBS can bidirectionally modulate neural activity: intermittent TBS (iTBS) typically facilitates cortical excitability, whereas continuous TBS (cTBS) suppresses it (Huang et al., 2005). Applying these protocols provides a direct method to manipulate the activity and observe its subsequent impact on throwing performance.

The purpose of the present study was to investigate the causal role of the SMA in stabilizing motor output during a dynamic throwing task in healthy novices. We employed iTBS and cTBS over the SMA and assessed the participants’ throwing performance in a maximal-effort throwing task, where signal-dependent noise is theoretically maximized.

Additionally, a submaximal throwing task was included to determine if SMA modulation affects the “sense of effort” as reflected in ball speed. To further explore the relationship between objective performance and conscious awareness, we also obtained introspective ratings of perceived speed and accuracy using visual analog scales (VAS). This allowed us to evaluate whether the SMA-mediated regulation of motor output operates as a subconscious process or is reflected in the subjective sense of motor performance. We hypothesized that if the SMA contributes to the stabilization of motor output under high-drive conditions, modulating its excitability would alter throwing performance. Specifically, we predicted that increasing the SMA excitability (via iTBS) would enhance the ability to control variability during maximal effort throwing, thereby mitigating the increased motor variability loss typically associated with high throwing speeds.

## Method

### Participants

Fourteen healthy, right-handed adults participated in the present study (mean ± standard deviation (SD), age: 23.0 ± 3.1 years old, height: 1.69 ± 0.07 m, weight: 61.6 ± 0.7 kg). The participants consisted of 4 females and 10 males, and none of them had prior competitive experience in baseball or softball. All participants reported no history of neurological or orthopedic disorders. All participants provided written informed consent. The experimental procedures were approved by the University of Tokyo’s local ethics committee (approval code: 942-5), and the study was conducted in accordance with the Declaration of Helsinki (1964).

### Ball-throwing task

All experiments were conducted indoors in a controlled laboratory environment. A standard rubber baseball (Type M; Kenko Co., Ltd., Tokyo, Japan) was used for the throwing task. The ball had a diameter of 72 mm, and a weight of approximately 138 g. Participants performed an overhand throwing task with their right hand, in which they threw a ball toward a target positioned 9 m away (target center: 1.1 m above the floor; target size: 0.2 × 0.2 m; Fig. 1A). The ball-throwing protocol consisted of two tasks: a submaximal throwing task (10 throws) and a maximal-effort throwing task (20 throws). Ten trials were performed in the submaximal task to evaluate the regulation of subjective motor drive (i.e., sense of effort), and thus, only ball speed was utilized as the outcome measure, whereas 20 trials were performed in the maximal task to comprehensively evaluate both throwing speed and ball arrival position. In the submaximal throwing task, participants were instructed to throw the ball toward the target using approximately 50 % of their subjective maximal effort. In the maximal throwing task, participants were instructed to “throw as fast and as accurately as possible toward the target.” In this maximal throwing task, introspective ratings of perceived throwing speed and accuracy were obtained on each trial using VAS. After throwing the ball, participants rated both the perceived speed and perceived accuracy of their throw using scales displayed on a 16-inch monitor. The VAS consisted of a 25-cm horizontal line anchored with “0” at the left end and “100” at the right end. For the perceived speed scale, 0 represented “could not throw fast at all” and 100 represented “could throw as fast as possible with maximal effort”. For the perceived accuracy scale, 0 represented “could not throw accurately at all” and 100 represented “could throw accurately”. The pixel coordinates of the selected points on the scale were converted into VAS scores ranging from 0 to 100 using numerical analysis software (MATLAB 2024b, The MathWorks Inc., US). The values were z-scored for each participant. Ball movements were recorded using a camera (DSC-RX10M4, SONY, Japan; 120 fps) placed on the participant’s back. Ball speed was recorded using a radar gun (Stalker Pro 2, Applied Concepts Inc. / Stalker Radar, US), which was fixed on a tripod and placed on the participant’s back (Fig. 1A).

**Fig. 1.**
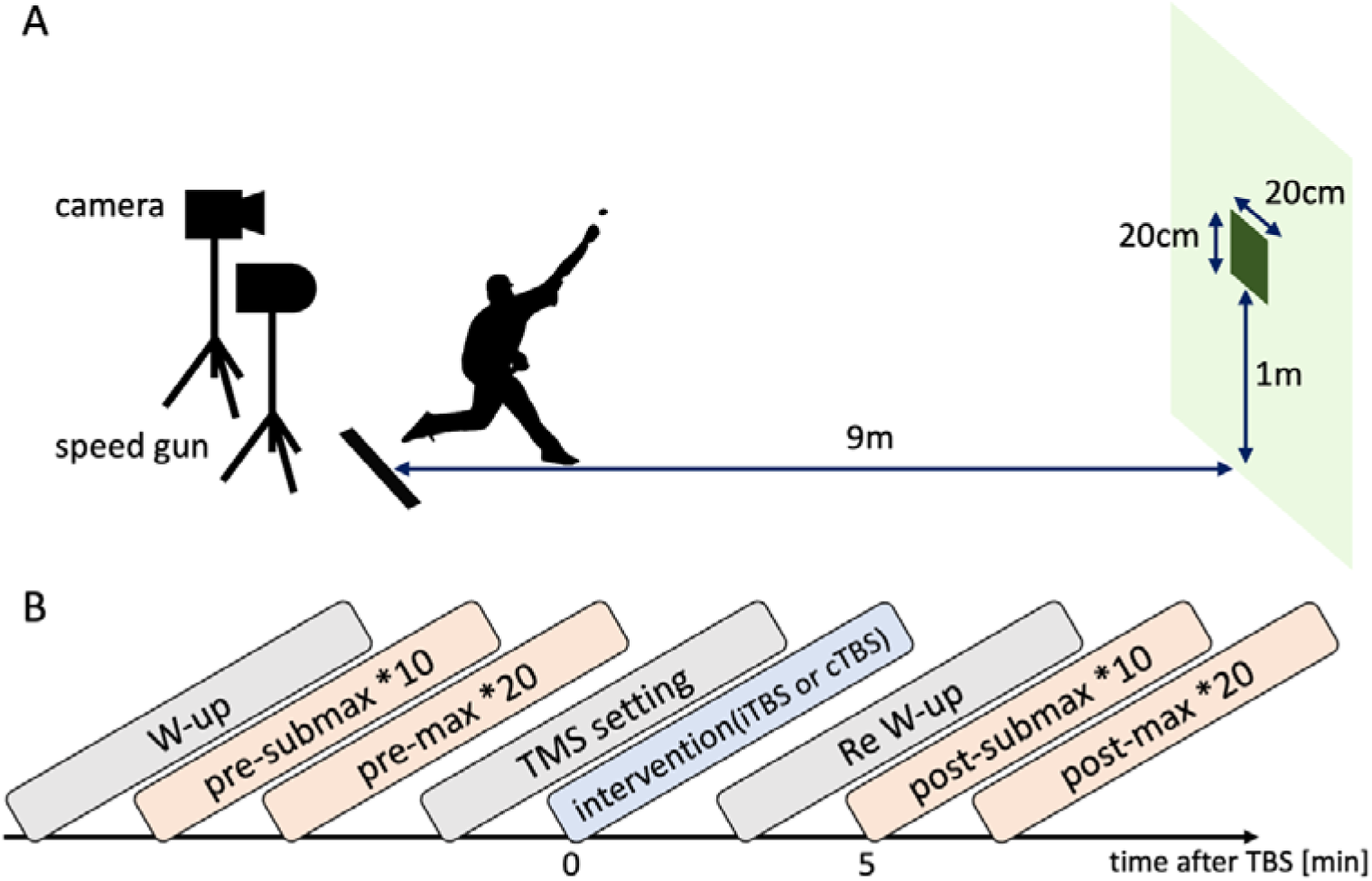
Overview of ball throwing task and experimental procedure. (A) Experimental setup for ball throwing task. A speed gun and a camera positioned behind the participants, were used to measure ball speed and arrival position, respectively. The target was positioned 9 m from the throwing position and 1.1 m above the floor. (B) Experimental timeline. The post-submaximal measurement was started at 5 minutes after TBS protocol.

### Theta burst stimulation (TBS)

Before administering the TBS protocols, we first identified the optimal stimulation site (i.e., hotspot) over the left primary motor cortex to determine the stimulation intensity. After cleansing the skin with alcohol, two bipolar Ag/AgCl surface electrodes (Vitrode F-150S, Nihon Kohden, Japan) were positioned on the right Flexor Carpi Radialis (FCR) muscle belly, with a grounding electrode near the right elbow. The FCR was selected as the target muscle because it acts as a prime mover for wrist flexion, playing a crucial role in the ball release phase of the overhand throwing motion (Aoyama et al., 2022; DiGiovine et al., 1992). TMS was applied at rest over the left primary motor cortex using a biphasic magnetic stimulator (DuoMAG XT-100, DEYMED Diagnostic s.r.o., Czech Republic) with a figure-of-eight coil (100 mm outer diameter; DEYMED Diagnostic s.r.o., Czech Republic). The hotspot was determined as the site where the largest MEP amplitude was elicited from the FCR muscle. A neuronavigation system (Brainsight, Rouge Research, Canada) ensured accurate placement over the hotspot. The coil was positioned tangentially to the scalp with the handle pointing backward and laterally at approximately 45° to the sagittal plane, inducing a posterior-anterior current in the cortex. The resting motor threshold (rMT) was defined as the lowest TMS intensity required to produce MEPs with peak-to-peak amplitudes greater than 50 μV in at least five out of ten successive trials while the muscle was fully relaxed (Rossini et al., 2015).

Each participant received two TBS protocols (iTBS and cTBS) administered with a biphasic magnetic stimulator (DuoMAG XT-100, DEYMED Diagnostic s.r.o., Czech Republic) with a figure-of-eight-shaped coil (100-mm outer diameter; DEYMED Diagnostic s.r.o., Czech Republic) in the separate experiment with random order. In brief, iTBS involved delivering three pulses at a frequency of 50 Hz per burst, applied every 200 ms for 2 seconds (totaling 600 pulses). For cTBS, a continuous 40-second burst stimulation was applied (600 pulses). The stimulation site, targeting the SMA, was set at the Montreal Neurological Institute (MNI) coordinates (-5, -51, 61) (Zénon et al., 2015) and designated using a neuronavigation system (Brainsight, Rouge Research, Canada). The coil was oriented to induce a posterior-anterior current. The target intensity of TBS was set at 80% of rMT.

### Experimental procedures

The experimental procedure is schematically illustrated in Fig. 1B. The present study utilized a crossover design. Participants completed two experimental sessions on separate days, with an interval of at least 7 days between sessions to prevent carry-over effects. The order of the stimulation conditions (iTBS or cTBS) was randomized and counterbalanced across participants. The experimental procedures were identical for both sessions, except for the type of stimulation applied. After completing a warm-up and practice session for the ball-throwing task, measurements and the TBS protocols began. Measurements were taken before (pre) and after (post) the TBS protocols. Each measurement session consisted of 10 submaximal throws followed by 20 maximal throws, and the order of these throwing tasks was fixed for all participants. After the pre-measurement, hotspot identification and TBS were conducted.

Following the TBS protocol, participants completed a light warm-up. The post-measurement began 5 minutes after the TBS protocol.

### Data and statistical analysis

The position coordinates of the pitch location were obtained using camera images and numerical analysis software (MATLAB 2024b, The MathWorks Inc., US). The points on the pitch in the camera images were obtained by digitizing the center point of the ball at the moment of arrival. The frames in which the ball arrived were selected by qualitative observation of the camera images. To obtain the position coordinates, we calibrated three points in the horizontal direction and two points in the vertical direction (at intervals of 2.9 m), giving a total of 6 calibration points for the transformation of the position coordinates. Pitch position coordinates were calculated using the 2D direct linear transformation (2DDLT) (Kusafuka et al., 2023; Shinya et al., 2017).

To evaluate the spatial throwing performance, we quantified two distinct components of error: variable error (precision) and constant error (accuracy). The two-dimensional distribution of pitch locations for each participant was fitted to a bivariate normal distribution. We calculated the area of the 95% confidence ellipse as an index of variable error(Hancock et al., 1995; Shinya et al., 2017). This metric reflects the precision of the throws, with a smaller area indicating tighter clustering of the pitch locations regardless of their proximity to the target. Additionally, we calculated the Euclidean distance between the center of the 95% confidence ellipse (i.e., the mean pitch location) and the center of the target as an index of constant error.

This metric represents the accuracy of the throws from the intended target.

Statistical analyses were performed using numerical analysis software (MATLAB 2024b, The MathWorks Inc., US). To evaluate the effects of the stimulation type and time on the measured variables (i.e., ball speed, ellipse area, distance to target, and introspective ratings), two-way repeated-measures analysis of variance (two-way rmANOVA) was conducted with within-subject factors of Stimulation (iTBS vs. cTBS) and Time (Pre vs. Post). When significant interaction effects were observed, post hoc comparisons were performed using paired t-tests with Bonferroni correction. Effect sizes were estimated using partial eta squared (η²_p_) for rmANOVA and Cohen’s d for t-tests. The significance level was set at p < 0.05.

## Result

Figure 2 shows the group means of the submaximal ball speed and maximal ball speed.

**Fig. 2.**
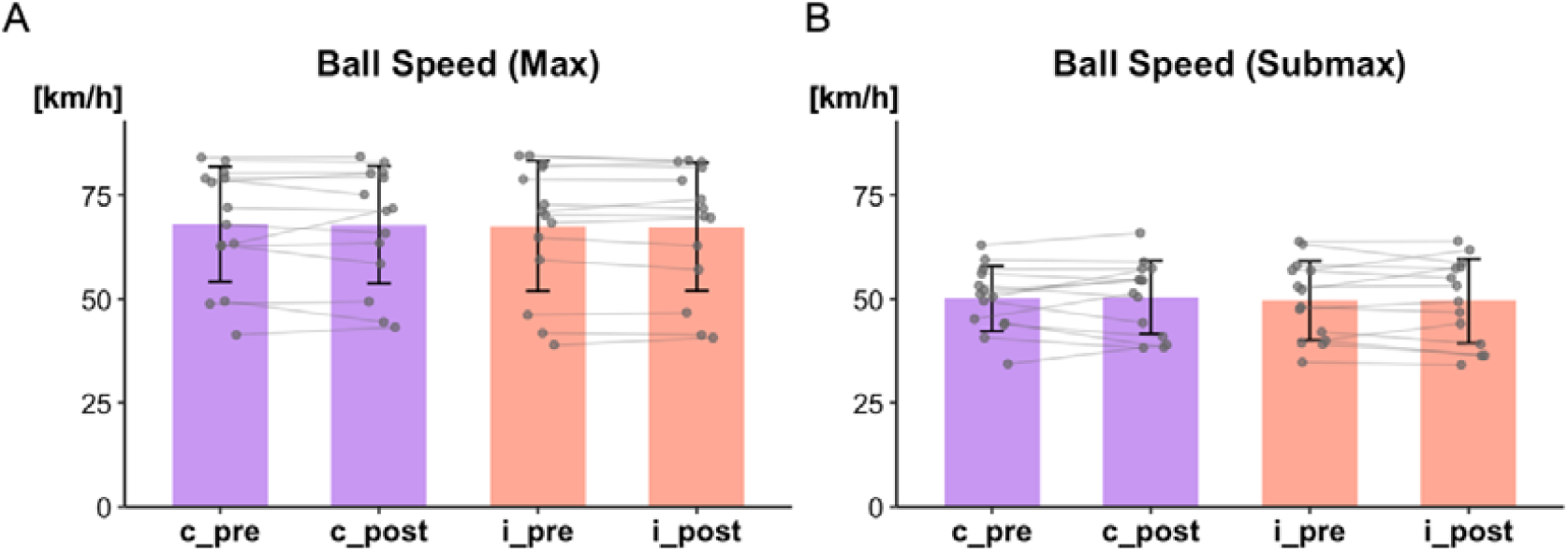
Changes in maximal ball speed (A) and submaximal ball speed (B) were assessed following cTBS and iTBS protocols. Bar plots show mean and error bar shows standard deviation (SD); gray circles represent individual participants.

For the submaximal ball speed, two-way rmANOVA did not show any significant main effects or interactions. [Stimulation: F (1, 13) = 0.198, p = 0.663, η²_p_ = 0.015, Time: F (1, 13) = 0.033, p = 0.858, η²_p_ = 0.002, interaction (Stimulation * Time): F (1, 13) = 0.126, p = 0.727, η²_p_ = 0.009]. For the maximal ball speed, two-way rmANOVA did not show any significant main effects or interactions. [Stimulation: F (1, 13) = 0.225, p = 0.642, η²_p_ = 0.017, Time: F (1, 13) = 0.008, p = 0.771, η²_p_ = 0.006, interaction (Stimulation * Time): F (1, 13) = 0.001, p = 0.972, η²_p_ < 0.001].

Figure 3 shows the group means of the area of the 95% confidence ellipse and the distance between the center of the 95% confidence ellipse and the target. For the area of the 95% confidence ellipse, two-way rmANOVA did not show any significant main effects. [Stimulation: F (1, 13) = 1.206, p = 0.292, η²_p_ =0.008, Time: F (1, 13) = 0.543, p = 0.474, η²_p_ =0.004], while a significant interaction was observed [Stimulation * Time: F (1, 13) = 10.099, p = 0.007, η²_p_ = 0.437]. Following the significant interaction, post hoc paired t-tests with Bonferroni correction revealed no significant difference between pre and post in the cTBS condition [t (13) = -1.16, p = 0.266, d = 0.208], whereas the iTBS condition showed a significant decrease from pre to post [t (13) = 2.56, p = 0.023, d = 0.539]. Comparisons between stimulation conditions indicated no significant difference between cTBS and iTBS at pre [t (13) = -1.24, p = 0.235, d = 0.168], while at post, the area was significantly smaller in the iTBS condition than in the cTBS condition [t (13) = 2.34, p = 0.035, d = 0.569]. For the distance between the center of the 95% confidence ellipse and the target, two-way rmANOVA did not show any significant main effects of time point or interactions [Time: F (1, 13) = 0.087, p = 0.771, η²_p_ = 0.006, interaction (Stimulation * Time): F (1, 13) = 1.195, p = 0.294, η²_p_ = 0.084] while a significant main effect of Stimulation was observed [F (1, 13) = 5.168, p = 0.040, η²_p_ = 0.284].

**Fig. 3.**
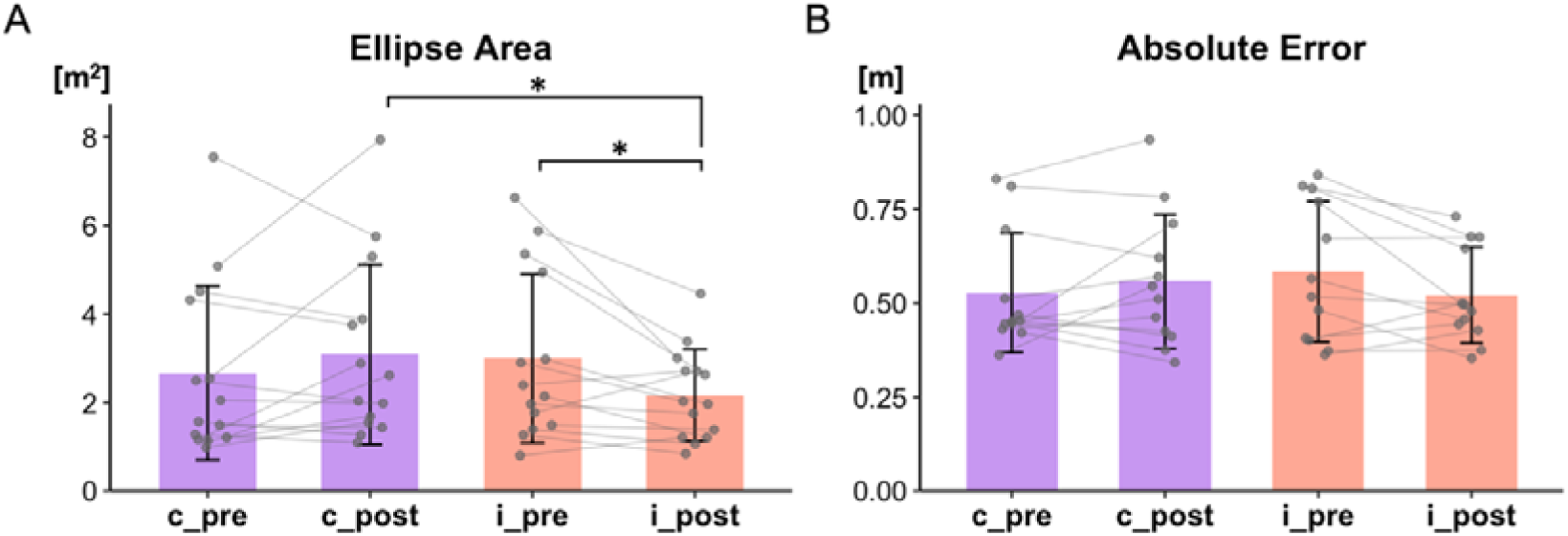
Changes in the area of the 95% confidence ellipse (A) and the distance between the center of the 95% confidence ellipse and the target (i.e., absolute error) (B) were assessed following cTBS and iTBS protocols. Bar plots show mean and error bar shows SD; gray circles represent individual participants. *p < 0.05 pre vs. post (Bonferroni-corrected).

Figure 4 shows the group means of the introspective rating (VAS) of speed and accuracy. For the VAS of speed, two-way rmANOVA did not show any significant main effects or interactions. [Stimulation: F (1, 13) = 0.012, p = 0.913, η²_p_ = 0.001, Time: F (1, 13) = 4.214, p = 0.064, η²_p_ = 0.277, interaction (Stimulation * Time): F (1, 13) = 0.002, p = 0.961, η²_p_ < 0.001]. For the VAS of accuracy, two-way rmANOVA did not show any significant main effects of Stimulation or interaction [Stimulation: F (1, 13) = 1.336, p = 0.268, η²_p_ = 0.093, interaction (Stimulation * Time): F (1, 13) = 0.198, p = 0.663, η²_p_ = 0.015], while a significant main effect of Time was observed [F (1, 13) = 7.096, p = 0.022, η²_p_ = 0.392]. Following the significant main effect of Time, post hoc paired t-test shows that the value of VAS (accuracy) was significantly greater at post rather than pre [t (13) = -2.664, p = 0.022, d = 1.538].

**Fig. 4.**
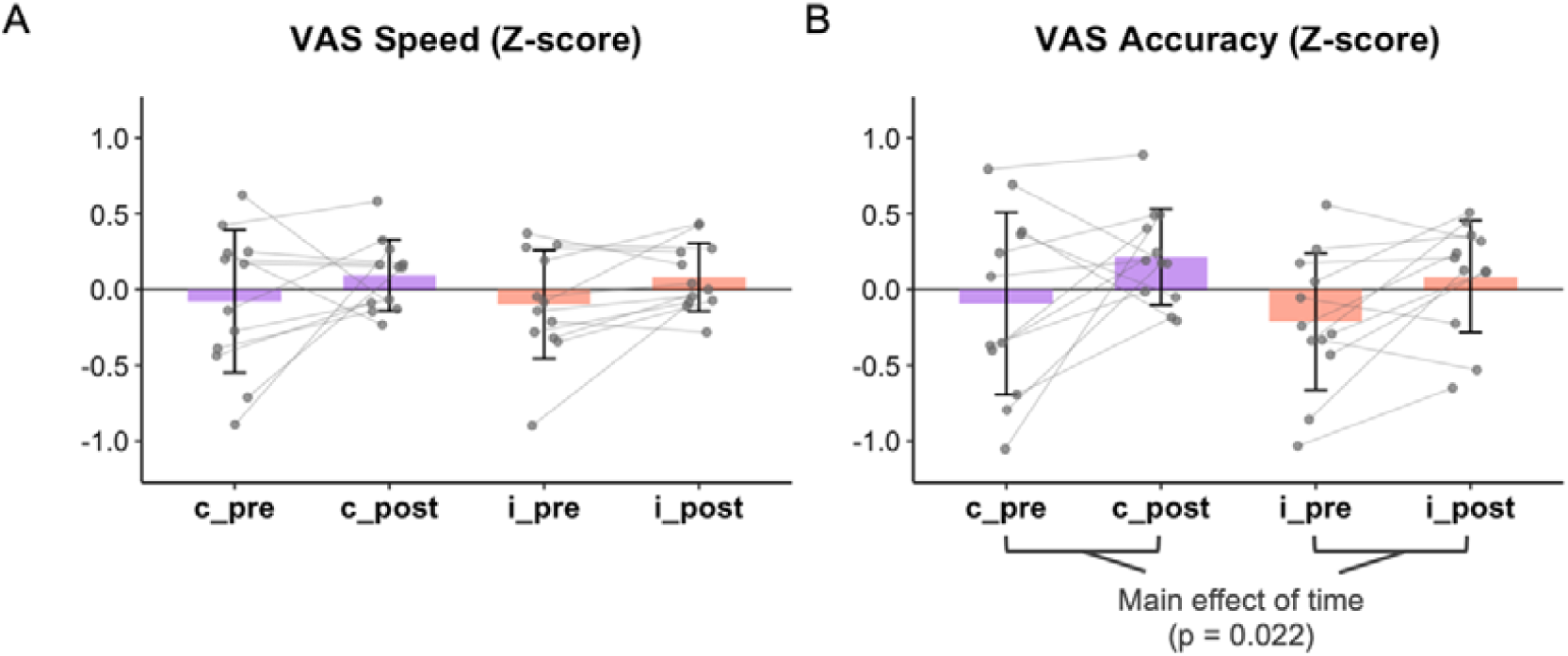
Changes introspective rating of speed (A) and accuracy (B) were assessed following cTBS and iTBS protocols. Data are z-transformed. Bar plots show mean and error bar shows SD; gray circles represent individual participants.

## Discussion

The primary aim of the present study was to elucidate the causal role of the SMA in stabilizing motor output during a dynamic throwing task. The most significant finding was that increasing SMA activity via iTBS significantly reduced the variable error of pitch location during maximal-effort throwing, without compromising ball speed. In contrast, inhibiting SMA activity via cTBS did not result in significant changes in throwing performance compared to the pre-stimulation baseline. Furthermore, neither stimulation protocol elicited significant effects during the submaximal throwing task. Collectively, these results support our hypothesis that the SMA plays a pivotal role in regulating motor variability, particularly under maximal effort conditions where signal-dependent noise is theoretically maximized.

The reduction in variable error under the iTBS condition can be interpreted through the SMA’s role in temporal coordination. Achieving both speed and accuracy in throwing requires the precise spatiotemporal coordination of multi-joint movements. Previous studies have indicated that the SMA is essential for maintaining temporal consistency (Jacobs et al., 2009) and facilitating anticipatory postural adjustments (Hasui et al., 2022). High-speed throwing requires rapid and synchronized muscle recruitment, a process highly susceptible to neural noise. By increasing the SMA excitability, iTBS may have reinforced the neural circuits responsible for timing accuracy and inter-joint coordination. Crucially, previous research has demonstrated that minimizing variability at the moment of ball release is a primary determinant of pitch location consistency (Kusafuka et al., 2020, 2022). Therefore, the enhanced SMA function likely constrained the temporal variability of the ball release parameters, leading to the observed improvement in endpoint precision even under high-speed conditions.

A particularly noteworthy aspect of our results is that the improvement in throwing performance was specific to precision (i.e., variable error), and this occurred without a concomitant decrease in ball speed. Although the classic motor control principle is broadly known as the speed-accuracy trade-off (Fitts, 1954), physiologically, high movement speeds tend to degrade performance primarily by amplifying motor variability due to signal-dependent noise (Faisal et al., 2008; Gandevia, 1987). The present results indicate that iTBS to the SMA can counteract the increase in variability typically associated with high throwing speeds, facilitating tighter spatial clustering while maintaining maximal speed. This pattern suggests that the observed improvement was not due to a strategic shift—such as sacrificing speed to improve target accuracy—but rather due to an enhancement in the efficiency of the motor control system, specifically in its capacity to constrain endpoint variability under high-speed conditions.

In contrast to the facilitatory effects of iTBS, cTBS did not result in a significant deterioration of throwing performance compared to the pre-stimulation baseline. One possible reason for this lack of effect may be the high baseline variability characteristic of novices. Because novices inherently exhibit high trial-to-trial variability in maximal effort throwing, any subtle performance deficits induced by the SMA suppression might have been difficult to distinguish from this intrinsic noise. This is consistent with the notion that the behavioral impact of non-invasive brain stimulation is state-dependent, where the observed effect is often influenced by the initial performance (Silvanto et al., 2008). In the case of novices, whose motor output is already inconsistent, the inhibitory impact of cTBS may be less apparent than it would be in a highly stable expert system. While these behavioral factors may explain the lack of significant decrement, the underlying neural processes remain to be fully elucidated. Future research should integrate neurophysiological measurements, such as Electroencephalography (EEG), to directly monitor cortical activity during the task (Miniussi & Thut, 2010; Ziemann, 2011). Such an approach would clarify whether the absence of a behavioral effect reflects a true lack of cortical modulation or a successful neural compensation within the motor network. Investigating these internal mechanisms will provide a more definitive understanding of how the SMA contributes to the control of complex, whole-body movements.

Interestingly, our findings revealed a complex dissociation between introspective ratings and objective motor performance, emphasizing that the SMA’s role in motor stabilization may operate independently of conscious awareness. We observed a significant global increase in the VAS ratings for accuracy over time; however, this subjective improvement was not reflected in the objective accuracy (constant error) of ball arrival position, which remained unchanged across all conditions. This suggests that the perceived improvement in “hitting the target” likely represents a psychological byproduct of task familiarization or a response bias—where repeated execution fosters a subjective sense of mastery despite no significant actual reduction in systematic error (Bjork et al., 2013; Simon & Bjork, 2001).

Furthermore, we cannot exclude the possibility of a placebo effect on introspection induced by the stimulation procedure itself. The expectation of receiving a neuromodulatory intervention may have created a biased perception of performance enhancement, independent of any actual change in motor output(Pollo et al., 2001). Crucially, while this subjective confidence of accuracy increased for all participants, a significant reduction in objective precision (variable error) of ball arrival position was exclusive to the iTBS condition. This indicates that participants were unable to consciously distinguish between a generalized sense of confidence and the specific physiological reduction in motor variability. The fact that iTBS selectively enhanced precision independently of subjective ratings suggests that the SMA-mediated noise reduction is a subconscious process (Charles et al., 2013; Fleming & Daw, 2017). In other words, while the “sense of accuracy” may be a metacognitive construct prone to practice effects, the stabilization of motor output appears to be a distinct neurophysiological process driven by the SMA’s capacity to constrain signal-dependent noise during high-speed movements (Blakemore et al., 2002; Nachev et al., 2008).

Despite the novel insights provided by this study, several limitations should be acknowledged. First, the participants were limited to novices without competitive baseball experience. While this allowed us to examine the fundamental role of the SMA in an unrefined motor system, it remains unclear whether these findings can be generalized to elite athletes, whose motor networks are highly optimized. In experts, the inhibitory impact of cTBS might be more apparent due to their stable motor output and minimal intrinsic noise. Unlike in novices, where inherent variability may mask subtle deficits, the SMA suppression in a highly tuned expert system could lead to significant performance degradation. Second, we targeted the SMA based on standardized MNI coordinates, not individualized MRI-based targeting. While this approach ensured consistent coil positioning across participants, it may not have adequately accounted for the anatomical variability among individuals, a limitation that is particularly relevant given the SMA’s relatively small cortical extent and could have reduced targeting precision.

In conclusion, the present study demonstrates that the SMA plays a causal role in stabilizing motor output during dynamic, high-speed movements. Facilitating the SMA excitability via iTBS selectively enhanced throwing precision in novices—reducing variable error without compromising ball speed—while suppressing SMA activity via cTBS yielded no significant performance changes. Crucially, the observed improvement in precision was dissociated from the participants’ introspective ratings, suggesting that the SMA-mediated noise reduction operates as a sub-perceptual process. These findings provide evidence that the SMA functions as a critical neural node for constraining signal-dependent noise under high-motor-drive conditions. Furthermore, our results highlight the potential of non-invasive brain stimulation as a tool for enhancing motor performance, offering a foundation for future applications in sports science and the optimization of complex motor skills.

## References

Aoyama, T., Ae, K., & Kohno, Y. (2022). Interindividual differences in upper limb muscle synergies during baseball throwing motion in male college baseball players. Journal of Biomechanics, 145, 111384. 10.1016/J.JBIOMECH.2022.111384

Bjork, R. A., Dunlosky, J., & Kornell, N. (2013). Self-regulated learning: Beliefs, techniques, and illusions. Annual Review of Psychology, 64(Volume 64, 2013), 417–444. 10.1146/ANNUREV-PSYCH-113011-143823/CITE/REFWORKS

Blakemore, S. J., Wolpert, D. M., & Frith, C. D. (2002). Abnormalities in the awareness of action. Trends in Cognitive Sciences, 6(6), 237–242. 10.1016/S1364-6613(02)01907-1

Cárdenas-Morales, L., Nowak, D. A., Kammer, T., Wolf, R. C., & Schönfeldt-Lecuona, C. (2010). Mechanisms and applications of theta-burst rTMS on the human motor cortex. Brain Topography, 22(4), 294–306. 10.1007/S10548-009-0084-7/TABLES/3

Charles, L., Van Opstal, F., Marti, S., & Dehaene, S. (2013). Distinct brain mechanisms for conscious versus subliminal error detection. NeuroImage, 73, 80–94. 10.1016/J.NEUROIMAGE.2013.01.054

DiGiovine, N. M., Jobe, F. W., Pink, M., & Perry, J. (1992). An electromyographic analysis of the upper extremity in pitching. Journal of Shoulder and Elbow Surgery, 1(1), 15–25. 10.1016/S1058-2746(09)80011-6

Faisal, A. A., Selen, L. P. J., & Wolpert, D. M. (2008). Noise in the nervous system. Nature Reviews Neuroscience 2008 9:4, 9(4), 292–303. 10.1038/nrn2258

Fitts, P. M. (1954). The information capacity of the human motor system in controlling the amplitude of movement. Journal of Experimental Psychology, 47(6), 381–391. 10.1037/H0055392

Fleming, S. M., & Daw, N. D. (2017). Self-Evaluation of Decision-Making: A General Bayesian Framework for Metacognitive Computation. Psychological Review, 124(1), 91. 10.1037/REV0000045

Gandevia, S. C. (1987). Roles for perceived voluntary motor commands in motor control. Trends in Neurosciences, 10(2), 81–85. 10.1016/0166-2236(87)90030-0

Hancock, G. R., Butler, M. S., & Fischman, M. G. (1995). On the Problem of Two-Dimensional Error Scores: Measures and Analyses of Accuracy, Bias, and Consistency. Journal of Motor Behavior, 27(3), 241–250. 10.1080/00222895.1995.9941714

Hasui, N., Mizuta, N., Taguchi, J., Nakatani, T., & Morioka, S. (2022). Effects of Transcranial Direct Current Stimulation over the Supplementary Motor Area Combined with Walking on the Intramuscular Coherence of the Tibialis Anterior in a Subacute Post-Stroke Patient: A Single-Case Study. Brain Sciences, 12(5). 10.3390/BRAINSCI12050540

Huang, Y. Z., Edwards, M. J., Rounis, E., Bhatia, K. P., & Rothwell, J. C. (2005). Theta burst stimulation of the human motor cortex. Neuron, 45(2), 201–206. 10.1016/j.neuron.2004.12.033

Iezzi, E., Suppa, A., Conte, A., Li Voti, P., Bologna, M., & Berardelli, A. (2011). Short-term and long-term plasticity interaction in human primary motor cortex. European Journal of Neuroscience, 33(10), 1908–1915. 10.1111/J.1460-9568.2011.07674.X;WGROUP:STRING:PUBLICATION

Jacobs, J. V., Lou, J. S., Kraakevik, J. A., & Horak, F. B. (2009). The supplementary motor area contributes to the timing of the anticipatory postural adjustment during step initiation in participants with and without Parkinson’s disease. Neuroscience, 164(2), 877–885. 10.1016/j.neuroscience.2009.08.002

Kusafuka, A., Kobayashi, H., Miki, T., Kuwata, M., Kudo, K., Nakazawa, K., & Wakao, S. (2020). Influence of Release Parameters on Pitch Location in Skilled Baseball Pitching. Frontiers in Sports and Active Living, 2. 10.3389/FSPOR.2020.00036/FULL

Kusafuka, A., Kudo, K., & Nakazawa, K. (2022). Control of Accuracy during Movements of High Speed: Implications from Baseball Pitching. Journal of Motor Behavior, 54(3), 304–315. 10.1080/00222895.2021.1960789

Kusafuka, A., Yamamoto, R., Okegawa, T., & Kudo, K. (2023). The ability to appropriately distinguish throws for different target positions. Frontiers in Sports and Active Living, 5, 1250938. 10.3389/FSPOR.2023.1250938/BIBTEX

Miniussi, C., & Thut, G. (2010). Combining TMS and EEG offers new prospects in cognitive neuroscience. Brain Topography, 22(4), 249–256. 10.1007/S10548-009-0083-8

Nachev, P., Kennard, C., & Husain, M. (2008). Functional role of the supplementary and pre-supplementary motor areas. Nature Reviews Neuroscience, 9(11), 856–869. 10.1038/NRN2478;KWRD

Okegawa, T., Yamasaki, D., Kaneko, N., & Nakazawa, K. (2025). Facilitation of supplementary motor area activity modulates the sense of effort: a theta burst stimulation study. Neuroscience Letters, 138482. 10.1016/J.NEULET.2025.138482

Picard, N., & Strick, P. L. (2001). Imaging the premotor areas. Current Opinion in Neurobiology, 11(6), 663–672. 10.1016/S0959-4388(01)00266-5

Pollo, A., Amanzio, M., Arslanian, A., Casadio, C., Maggi, G., & Benedetti, F. (2001). Response expectancies in placebo analgesia and their clinical relevance. Pain, 93(1), 77–84. 10.1016/S0304-3959(01)00296-2

Roach, N. T., Venkadesan, M., Rainbow, M. J., & Lieberman, D. E. (2013). Elastic energy storage in the shoulder and the evolution of high-speed throwing in Homo. Nature, 498(7455), 483–486. 10.1038/NATURE12267

Rossini, P. M., Burke, D., Chen, R., Cohen, L. G., Daskalakis, Z., Di Iorio, R., Di Lazzaro, V., Ferreri, F., Fitzgerald, P. B., George, M. S., Hallett, M., Lefaucheur, J. P., Langguth, B., Matsumoto, H., Miniussi, C., Nitsche, M. A., Pascual-Leone, A., Paulus, W., Rossi, S., … Ziemann, U. (2015). Non-invasive electrical and magnetic stimulation of the brain, spinal cord, roots and peripheral nerves: Basic principles and procedures for routine clinical and research application: An updated report from an I.F.C.N. Committee. Clinical Neurophysiology, 126(6), 1071–1107. 10.1016/j.clinph.2015.02.001

Saito, H., Kusafuka, A., Okegawa, T., Takao, S., Kaneko, N., Yokoyama, H., Takiyama, K., Takaki, K., & Nakazawa, K. (2025). Developmental changes in upper limb muscle synergies during throwing: A comparison between preschoolers and schoolers. IScience, 28(10), 113497. 10.1016/j.isci.2025.113497

Schmidt, R. A., &, et al. (1979). Motor-output variability: A theory for the accuracy of rapid motor acts. Psychological Review, 86(5), 415–451. 10.1037/0033-295X.86.5.415

Shinya, M., Tsuchiya, S., Yamada, Y., Nakazawa, K., Kudo, K., & Oda, S. (2017). Pitching form determines probabilistic structure of errors in pitch location. Journal of Sports Sciences, 35(21), 2142–2147. 10.1080/02640414.2016.1258484

Silvanto, J., Muggleton, N., & Walsh, V. (2008). State-dependency in brain stimulation studies of perception and cognition. Trends in Cognitive Sciences, 12(12), 447–454. 10.1016/j.tics.2008.09.004

Simon, D. A., & Bjork, R. A. (2001). Metacognition in Motor Learning. Journal of Experimental Psychology: Learning Memory and Cognition, 27(4), 907–912. 10.1037/0278-7393.27.4.907

Understanding Motor Development: Infants, Children, Adolescents, Adults … - Jacqueline D Goodway, John C Ozmun, David L Gallahue - Google Books. (n.d.). Retrieved December 4, 2025, from https://books.google.co.jp/books?hl=en&lr=&id=h5KwDwAAQBAJ&oi=fnd&pg=PP1&ots=UaFQgMuwfv&sig=9GUFwSVP8-A3UDfxWoShG54xNOs&redir_esc=y#v=onepage&q&f=false

Van Den Tillaar, R., & Ettema, G. (2006). A comparison between novices and experts of the velocity-accuracy trade-off in overarm throwing. Perceptual and Motor Skills, 103(2), 503–514. 10.2466/PMS.103.2.503-514;WGROUP:STRING:PUBLICATION

Zénon, A., Sidibé, M., & Olivier, E. (2015). Disrupting the Supplementary Motor Area Makes Physical Effort Appear Less Effortful. Journal of Neuroscience, 35(23), 8737–8744. 10.1523/JNEUROSCI.3789-14.2015

Ziemann, U. (2011). Transcranial magnetic stimulation at the interface with other techniques: A powerful tool for studying the human cortex. Neuroscientist, 17(4), 368–381. 10.1177/1073858410390225;WGROUP:STRING:PUBLICATION

